# Phenome-wide and Genome-wide Analyses of Quality of Life in Schizophrenia

**DOI:** 10.1101/744045

**Authors:** Raha Pazoki, Bochao Danae Lin, Kristel R. van Eijk, Dick Schijven, Sinan Guloksuz, GROUP investigators, Jurjen J. Luykx

## Abstract

**Background:** Schizophrenia negatively impacts quality of life (QoL). A handful of variables from small studies have been reported to influence QoL of schizophrenia patients, but a study comprehensively dissecting the genetic and non-genetic contributing factors to QoL in these patients is currently lacking. We adopted a hypothesis-generating approach to assess the phenotypic and genotypic determinants of QoL in schizophrenia.

**Method:** The study population consisted of 1,119 patients with a psychotic disorder, 1,979 relatives and 586 healthy controls. Using linear regression, we tested >100 independent demographic, cognitive and clinical phenotypes for their association with QoL in patients. We then performed genome-wide association analyses of QoL and examined the association between polygenic risk scores (PRSs) for schizophrenia, major depressive disorder (MDD), and subjective wellbeing (SW) with QoL.

**Results:** We found nine phenotypes to be significantly and independently associated with QoL in patients, the most significant ones being negative (Beta=-1.17; SE=0.05, P=1×10^-83^; r^2^=53%), depressive (Beta=-1.07; SE=0.05; P=2×10^-79^; r^2^=51%) and emotional distress (Beta=-0.09; SE=0.01; P=4×10^-59^, r^2^=38%) symptoms. Schizophrenia and subjective wellbeing PRSs using various P-value thresholds were significantly and consistently associated with QoL (lowest association p-value = 6.8×10^-6^). Several sensitivity analyses confirmed the results.

**Conclusions:** Various clinical phenotypes of schizophrenia as well as schizophrenia and subjective wellbeing polygenic risk scores are associated with QoL in schizophrenia patients and their relatives. These may be targeted by clinicians to more easily identify vulnerable schizophrenia patients for further social and clinical interventions to improve their QoL.

## Introduction

Schizophrenia (SCZ) patients often experience adverse outcomes such as unemployment, frequent hospital admissions, long-term dependency on health care, and suicide. Premature mortality of patients with SCZ has been reported to be 3.5 times (Olfson et al., 2015) greater than that of adults in the general population. The societal costs of SCZ during a 12-month period have been estimated to be as high as $890 million in the United States (Evensen et al., 2015). All domains of quality of life (QoL; physical, psychological, and social) are severely decreased in SCZ compared to healthy controls. QoL is also increasingly becoming an important index for effectiveness of treatment in SCZ (Kane et al., 2016). Several variables have been shown to be associated with QoL among SCZ patients, e.g. age, gender, employment status, marital status, duration of illness, body mass index, antipsychotic medication, number of hospitalizations, knowledge level about schizophrenia, schizophrenia symptoms, coping mechanisms, and comorbid depression (Hasan and Tumah, 2019, Hofer et al., 2017, Hou et al., 2016, Karow et al., 2014, Rayan and Obiedate, 2017, Rotstein et al., 2018, Savill et al., 2016, Wang et al., 2017, Yamauchi et al., 2008). However, a comprehensive large-scale study using in-depth phenotyping to investigate factors associated with QoL in SCZ in a hypothesis-generating fashion is lacking. In addition, to the best of our knowledge, the genetic underpinnings of QoL have not been investigated. Recently published genome-wide association studies (GWASs) for SCZ (Ripke et al., 2014) and related traits such as major depressive disorder (MDD) (Wray et al., 2018) and subjective wellbeing (Okbay et al., 2016) provide a timely opportunity to investigate whether genetic mechanisms are at the root of QoL. Knowledge of clinical and genetic contributing factors to QoL in SCZ could inform clinicians to help identify vulnerable patients and optimize secondary preventive care and thus reduce burden of disease. This could be achieved through optimization of treatment regimens (e.g. psychosocial interventions or optimizing psychopharmacological treatments) and targeting clinical variables negatively influencing QoL. On a similar note, insight into genetic factors contributing to QoL could contribute to the early identification of vulnerable patients and in the future improve their outcome.

Here, we used a hypothesis-generating approach and investigated over 100 phenotypes to investigate factors related to QoL among SCZ patients. We additionally performed genetic risk scoring in patients, relatives and healthy controls to uncover associations between genetic susceptibility to SCZ, MDD and subjective wellbeing on the one hand and QoL on the other.

## Method

### Subjects and study design

All procedures contributing to this work comply with the ethical standards of the relevant national and institutional committees on human experimentation and with the Helsinki Declaration of 1975, as revised in 2008. All procedures involving human subjects/patients were approved by the medical-ethical committee of University Medical Center Utrecht (UMCU). All subjects provided written informed consent for the current study. The current study was performed within a cohort of 3,684 individuals including 1,119 SCZ patients, 1,059 siblings, 920 parents, and 586 controls (**Supplementary Results, Suppl. Figure 1**). We used two main subsets. The first subset included patients only (n=1,119) to test non-genetic contributing factors to QoL among SCZ patients. We chose this subset as we were interested in phenotypic contributing factors to QoL in patients, while other, larger cohort studies may be more appropriate to probe contributing factors to QoL in the general population. The second subset included patients, relatives and controls with genetic data available (n=2,265) to test genetic contributing factors to QoL. We chose this subset to increase statistical power and as intuitively genetic contributing factors to QoL may (partly) overlap between patients, relatives and controls. All participants were included from the Genetic Risk and Outcome of Psychosis (GROUP) study (Korver et al., 2012), a multi-center and large longitudinal study in the Netherlands and Belgium, investigating various psychological and genetic variables among SCZ patients and their relatives. The study population was followed up since 2004 in several mental health care institutions, both in the Netherlands and Belgium. The detailed phenotypic information of GROUP participants offers a unique and enriched database (Korver et al., 2012). Psychosis-related and demographic variables that were included in the analysis are presented in **Appendix 1 (Supplemental Methods)**. These variables may be divided into symptoms and experiences that were assessed with a range of (semi)-structured scales (including drug use); family loading for psychiatric disorders; social cognition; demographic variables; IQ; medication use data; and theory of mind scales. For the purpose of the current study, we only used the baseline assessments of the GROUP study (release 5.00) as we were interested in factors contributing to QoL in SCZ apparent in its early disease stages (the first psychotic episode of the GROUP participating patients had to occur within 10 years before this first assessment). **Supplementary Figure 1** shows a breakdown of the study sample.

### Quality of Life Assessment

Quality of life was assessed with the World Health Organization WHOQOL-BREF, an abbreviated version of the WHOQOL (World Health Organization Quality of Life scale). The self-report WHOQOL-BREF has been validated for a Dutch speaking population of psychiatric patients (Korver et al., 2012, Trompenaars et al., 2005, Gobbens and van Assen, 2016). Details of the Dutch WHOQOL-BREF are described elsewhere (Gobbens and van Assen, 2016). Details of our method to extract a single principal component derived from this questionnaire that was used for further analysis are described in the supplemental methods.

### Phenome-wide analyses

For the agnostic association analysis of QoL with the above explained demographic and clinical phenotypes (**Supplemental methods, Appendix 1**), we used data from patients with complete data on the QoL principal component, age, sex, and study site (N=925; **Supplementary Figure 1**). The number of independent variables was calculated by testing two-by-two correlations between variables using non-parametric Spearman correlation. Variables with correlation estimates > 0.3 and statistically significant correlations (P value< 0.05) were considered interdependent variables. This analysis resulted in 105 independent variables. Generalized linear models (GLM) adjusted for age, sex, and study site were then used to test the association of QoL with each of the clinical phenotypes. The statistical significance threshold for this association analysis was corrected for multiple testing using the Bonferroni-correction method, i.e. 4.76×10^-04^ (0.05 adjusted for 105 independent tests)(Shaffer, 1995). The variance in QoL explained by the phenotypes was calculated using R square values obtained from GLM. To then identify a set of variables that were associated with QoL independent of one another, we used the phenotypes that were associated with QoL at P<4.76×10^-04^ and selected the independent variables in a backward stepwise regression model. As a sensitivity analysis, we then regressed the most significantly associated phenotypes with ordinal estimates of QoL as opposed to the first principal component of QoL.

### Polygenic risk score analyses of QoL

Details of genotyping and GWAS of QoL can be found in the Supplementary methods. In brief, genotype data for 2,812 GROUP participants was generated on a customized Illumina IPMCN array with 570,038 single nucleotide polymorphisms (SNPs). Quality control procedures were performed using PLINK v1.9(Purcell et al., 2007). In total, 2,505 individuals and 275,021 SNPs passed these abovementioned QC steps. After merging with the phenotype file, 2,265 individuals were left for genetic analyses (**Supplemental Results, Supplementary Figure 1**).

Additional SNPs were imputed on the Michigan server (Das et al., 2016) using the HRC r1.1 2016 reference panel. Although likely underpowered, for the benefit of possible future meta-analyses and as a first exploratory approach we performed linear mixed models (LMM) association testing implemented in BOLT-LMM (v2.3) software(Loh et al., 2015) to assess associations between SNPs and QoL (**Supplemental methods**). BOLT-LMM corrects for confounding from population structure and cryptic relatedness. We used the generally accepted association P-value threshold of P <5×10^-8^ for genome-wide significance. We report those findings in the **Supplemental Results (Figures 7&8)**.

We used recent GWASs of SCZ (Ripke et al., 2014), MDD (Wray et al., 2018), and subjective wellbeing (Okbay et al., 2016) for PRS calculations (Choi et al., 2018). We chose the polygenic risk scores of these disorders as they are strongly associated with QoL in the general population (IsHak et al., 2015, Skevington and Böhnke, 2018, Camfield and Skevington, 2008, Domenech et al., 2018). To verify that PRS of other traits were indeed unlikely to be associated with QoL, the genetic correlations between the primary BOLT-LMM GWAS summary statistics and over 700 other disease traits were estimated using LD score regression (http://ldsc.broadinstitute.org/) (Zheng et al., 2017). As a quality control for PRS calculation, the SNPs that overlapped between the summary statistics GWASs (training datasets) and our dataset were extracted. Then, insertions or deletions, ambiguous SNPs, SNPs with minor allele frequency (MAF) <0.01 and imputation quality (R^2^) < 0.8 in both training and target datasets were excluded. To account for complicated LD structure of SNPs in the genome, these SNPs were clumped in two rounds using PLINK 1.90b3z (Chang et al., 2015) according to previously established methods (McLaughlin et al., 2017b, Schur et al., 2019); round 1 with the default parameters (physical distance threshold 250kb and LD threshold (R^2^) 0.5); round 2 with a physical distance threshold of 5,000kb and LD threshold (R^2^) 0.2. Additionally, we excluded all SNPs in genomic regions with strong or complex LD structures (e.g. the MHC region on chromosome 6; **Supplemental Results, Supplementary Table 1**). If only odds ratios (ORs) were reported in the summary statistics, ORs were log-converted to beta values as effect sizes. To prevent possible study population overlap impacting our results, all Dutch and Belgian individuals had been excluded from the SCZ GWAS (Ripke et al., 2014) to allow unbiased PRS computation (McLaughlin et al., 2017a). Sample overlap between GROUP data with MDD and subjective wellbeing GWAS samples is unlikely since all samples belong to different cohorts. To reassure that there was indeed minimal to no sample overlap between GROUP vs. MDD and subjective well-being samples, we checked the intercepts of the genetic covariances from LD score regression analyses between the GROUP GWAS vs. MDD and subjective well-being. Presence of sample overlap modifies this intercepts from zero (Bulik-Sullivan et al., 2015), while in our study all intercepts turned out to be close to zero (**Supplemental Methods & Supplemental Results, Supplementary table 2**). We constructed PRSs based on SCZ risk alleles weighted by their SCZ increasing effect estimate using the Purcell et al. method (Purcell et al., 2007, Purcell et al., 2009), i.e. using PLINK’s score function for 12 GWAS p-value thresholds: 5 × 10^-8^, 5 × 10^-7^, 5 × 10^-6^, 5 × 10^-5^, 5 × 10^-4^, 5 × 10^-3^, 0.05, 0.1, 0.2, 0.3, 0.4 and 0.5. PRSs were calculated for 2,505 patients, relatives and controls (those remaining after QC). Genetic data and QoL variables were available for patients, relatives, and controls. We thus performed statistical analyses for the association of PRSs with QoL in the whole sample after QC (N=2,265; **Supplemental Results, Supplementary Figure 1**) including patients, controls and family members. This approach provided the opportunity to investigate genetic susceptibility of these PRSs on QoL regardless of presence or absence of the disease. To claim significance for association analyses between PRS and QoL, we Bonferroni corrected the P-value for multiple testing (0.05/3=0.016).

### Data availability

All authors have continuous access to the data, both phenotypic and genotypic, collected in this study.

## Results

**Table 1** shows baseline characteristics of patients, relatives and controls. In the generalized linear model (GLM), 18 distinct variables were associated with QoL at the Bonferroni significance threshold of 4.76×10^-04^ in schizophrenia patients (**Figure 1, Table 2 & Supplemental Results, Supplementary Figure 3**). The statistically most significant phenotypes were negative (Beta=-1.17; SE=0.05, P=1×10^-83^; r^2^ model=53%), depressive (Beta=-1.07; SE=0.05; P=2×10^-79^; r^2^ model=51%), emotional distress (Beta=-0.09; SE=0.01; P=4×10^-59^, r^2^ model=38%), and general psychopathology (Beta=0.81; SE=0.06; P=3×10^-40^; r^2^ model=29%) symptoms (**Table 2**). In our regression model including these 18 variables, nine remained independently associated with QoL (p-value<0.05), explaining 58.55% of the variance in QoL. Ordered by decreasing level of significance these are: negative symptoms, global assessment of functioning, emotional distress, depressive symptoms, positive symptoms, remission status, cannabis craving, number of unmet needs, and excitement (**Table 2 & Supplemental Results, Supplementary Table 3)**. In addition, there was a negative age effect (Beta=-0.01; SE=0.003; P=3×10^-3^) on QoL in the backward stepwise model (**Supplemental Results, Supplementary Table 3**). Association analysis between ordinal estimates of QoL showed similar results (**Supplemental Results, Supplementary Figure 4**).

**Table 1.** Baseline characteristics for patients, siblings and controls.

| **Characteristics** | **Patients**<br>**(n=1,119)** | **Controls**<br>**(n=586)** | **Siblings**<br>**(n=1,059)** | **Parents**<br>**(n=920)** |
| --- | --- | --- | --- | --- |
| -Age in years, mean (sd) | 27.6(7.9) | 30.4(10.6) | 27.8(8.3) | 54.7(6.7) |
| -Gender, n (%) women | 267(23.9) | 317(54.1) | 577(54.5) | 528(57.4) |
| -IQ, estimated, mean (sd) | 95(16.1) | 109.7(15.1) | 102.8(15.6) | 103(17.0) |
| -Married/living together, n (%) | 97(9.3) | 234(41.1) | 411(40.2) | 153(70.8) |
| -Years of education , mean (sd) | 4 (2.1) | 5.4(1.8) | 5.1(2.1) | 5.1(2.3) |
| -Nicotine use, mean number of cigarettes daily (sd) | 11.7(11) | 3(6.5) | 4.9(8.4) | 4.3(8.8) |
| -Alcohol use, mean number of drinks per week (sd) | 6.6(12.1) | 6.1(8.5) | 6.4(8.6) | 8.1(10.6) |
| -Current use of Antipsychotics, % | 1062(95) | 0 (0) | 0 (0) | 2(0.22) |
| -Duration of Illness (years), mean (sd) | 4.2(4) | N/A | N/A | N/A |

**Table 2.** The 18 distinct clinical variables associated with QoL in the generalized linear model (N= 925 schizophrenia patients).

| **Variable** | **Scale** | **Standardized**<br>**Effect**<br>**Estimate** | **Standard**<br>**Error** | **Explained**<br>**Variance** | **P**<br>**Value** |
| --- | --- | --- | --- | --- | --- |
| **Negative symptoms, points** | **CAPE** | **-1.17** | **0.05** | **0.53** | **$1 \times 10^{-83}$** |
| **Depressive symptoms, points** | **CAPE** | **-1.07** | **0.05** | **0.51** | **$2 \times 10^{-79}$** |
| **Emotional distress, points** | **PANSS** | **-0.09** | **0.01** | **0.38** | **$4 \times 10^{-59}$** |
| **General psychopathology**<br>**symptoms, points** | PANSS | -0.81 | 0.06 | 0.29 | $3 \times 10^{-40}$ |
| **Global assessment of functioning**<br>**(disabilities), points\*** | **GAF** | **0.03** | **0** | **0.26** | **$2 \times 10^{-34}$** |
| **Positive symptoms, points** | **CAPE** | **-0.05** | **0** | **0.2** | **$1 \times 10^{-23}$** |
| **Number of unmet needs, points** | **CAN** | **-0.11** | **0.01** | **0.19** | **$3 \times 10^{-23}$** |
| **Remission status, yes** | **PANSS** | **-0.56** | **0.07** | **0.15** | **$8 \times 10^{-17}$** |
| **Excitement, points** | **PANSS** | **-0.06** | **0.01** | **0.13** | **$9 \times 10^{-13}$** |
| PAS total score | PANSS | -0.25 | 0.04 | 0.10 | $5 \times 10^{-11}$ |
| Proportion unmet needs | CAN | -0.69 | 0.11 | 0.10 | $2 \times 10^{-9}$ |
| Disorganization, points | PANSS | -0.03 | 0.01 | 0.11 | $3 \times 10^{-9}$ |
| Obsessive compulsive symptoms,<br>yes | Y-BOCS | -0.43 | 0.08 | 0.09 | $4 \times 10^{-8}$ |
| Suicidal attempts (lifetime), yes | Composite<br>file** | -0.43 | 0.08 | 0.09 | $6 \times 10^{-8}$ |
| **Cannabis craving, yes** | **OC-DUS** | **-0.31** | **0.06** | **0.14** | **$1 \times 10^{-07}$** |
| OCT Total score | OC-DUS | -0.28 | 0.06 | 0.13 | $1 \times 10^{-06}$ |
| Deficit syndrome | SDS | -0.17 | 0.04 | 0.06 | $7 \times 10^{-05}$ |
| Acathisia | BARS | -0.13 | 0.04 | 0.06 | $3 \times 10^{-04}$ |

CAPE: Community Assessment of Psychic Experiences; PANSS: positive and negative syndrome scale; GAF, Global assessment of functioning; CAN, The Camberwell Assessment of Need; Y-BOCS: Yale-Brown obsessive compulsive scale; OC-DUS: obsessive-compulsive drug use scale; SDS: Schedule for the Deficit Syndrome; BARS: Barnes Akathisia Rating Scale Global, a clinical assessment scale for acathisia \* Greater score indicates better functioning. \*\* This scale contains a range of questions probing health. Note: **the clinical variables in bold were independently associated with QoL in our stepwise regression model.**

**Figure 1.**
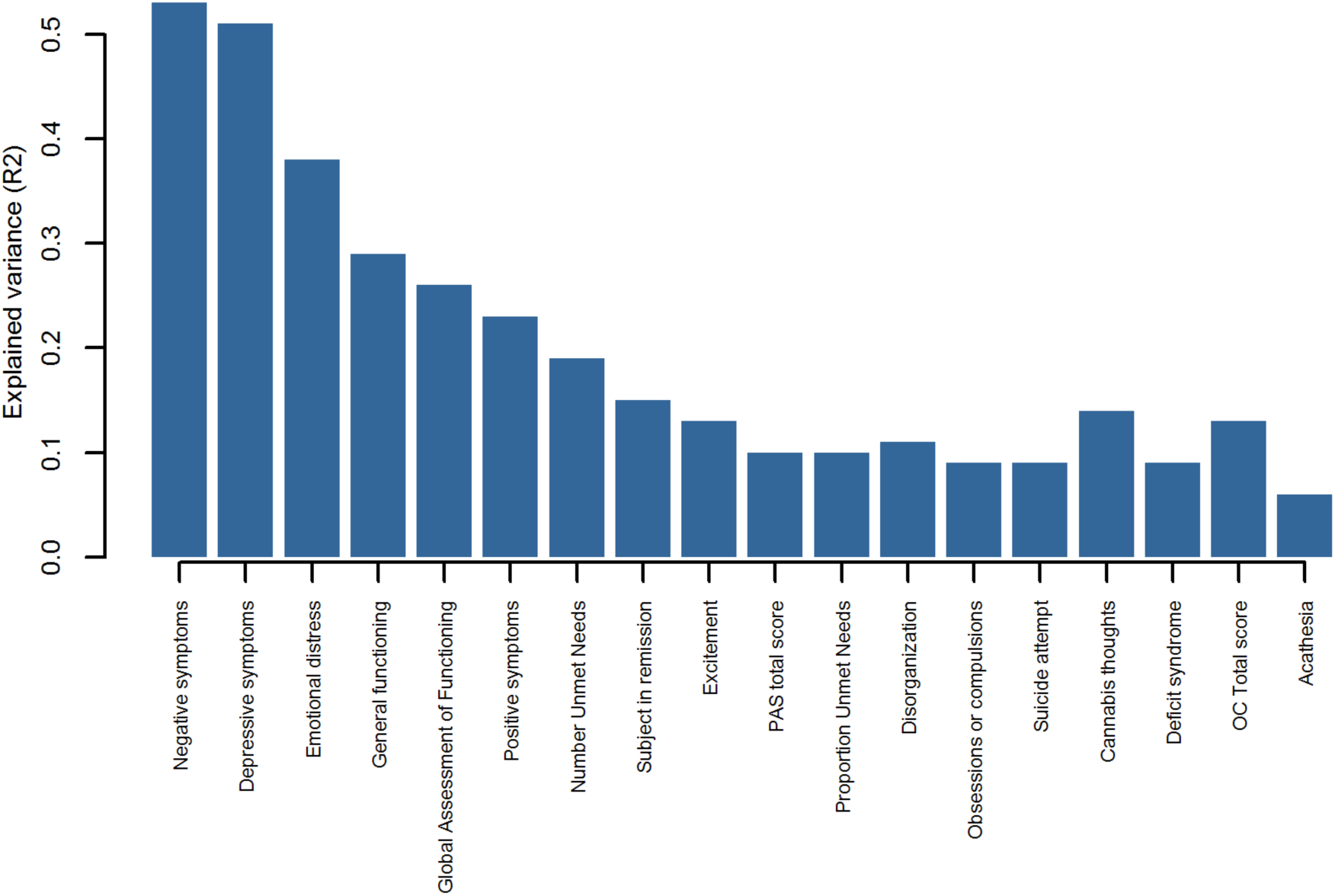
Results of the hypothesis-generating association analysis between clinical variables and QoL among SCZ patients with explained variance for QoL. Number unmet needs, measured using the CAN, The Camberwell Assessment of Need. Subject in remission: measured using the PANSS subject in remission tool. PAS: Personality Assessment Screener. Suicide attempt: assessed using the composite file (a questionnaire with closed questions designed for the GROUP study). Cannabis thoughts: thoughts about cannabis use, measured using the OC-DUS (obsessive compulsive drug use scale). Deficit syndrome: measured using the SDS (Schedule for the Deficit Syndrome). OC total score: obsessive compulsive symptoms score measured with the Yale-Brown obsessive compulsive scale. Akathisia, measured using the Barnes akathisia rating scale.

The variance in QoL explained by various PRSs (N=2,265) were 1.37% for SCZ, 1.37% for subjective wellbeing (**Figure 2)**, and 1.40% for MDD (**Supplemental Results, Supplementary Figure 5**) when using only genome-wide significant SNPs (P-value threshold (Pt) of 5×10^-8^). The most significant associations between PRS and QoL were observed for SCZ (Pt_0.5_; explained variance=1.58%, P =7×10^-6^; **Figure 1)**, subjective wellbeing (Pt_0.4_; explained variance=1.82%, P=0.004; **Figure 1**), and MDD (Pt_0.005_; explained variance=1.62%, P= 0.01; **Supplemental Results, Supplementary Figure 5**). As a sensitivity analysis, we repeated the SCZ PRS analysis on patients only (N=633) and confirmed the same pattern of association with QoL and the same Pt of 0.5 showing most significant association results (**Supplemental Results, Supplementary Figure 6).** As expected given relatively low statistical power, genetic correlation analysis in LD Hub showed no statistically significant results. Confirming our rationale for investigating the PRSs chosen in the current study, genetic correlations of Qol with SCZ (Ripke et al., 2014) and subjective wellbeing (Okbay et al., 2016) were the strongest, in the expected direction (**Supplemental Results, Supplementary Table 2**).

**Figure 2.**
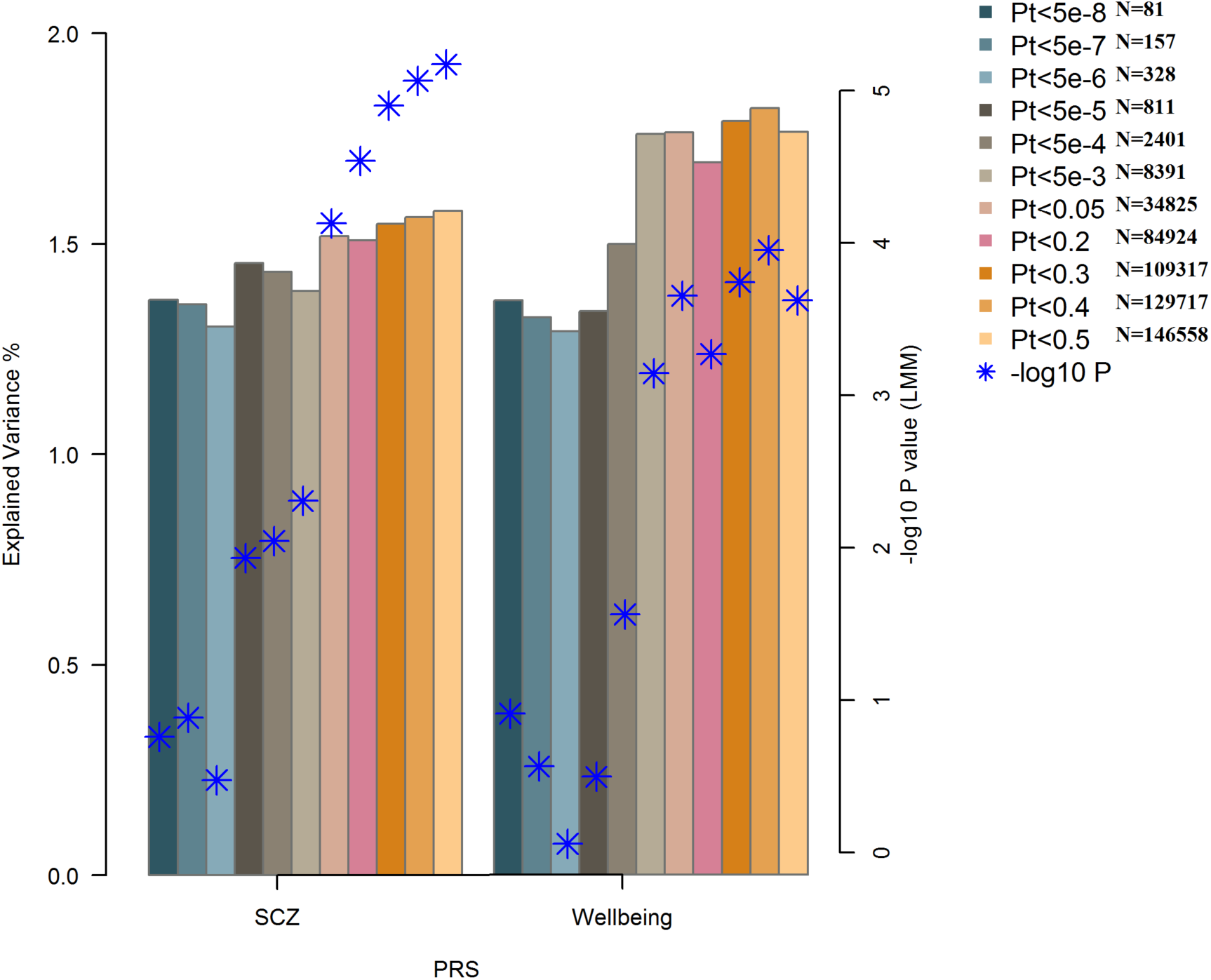
Bar plot illustrating explained variance for association of polygenic risk scores of SCZ (schizophrenia) and subjective wellbeing with QoL (quality of life). The figure illustrates the results using linear mixed models. Displayed are the number of SNPs (N), the strengths of the association results (-log 10 P) and explained variances per Pt (p-value threshold).

SCZ Pt_0.5_ (P =7×10^-6^), MDD Pt_0.005_ (P= 0.01), and subjective wellbeing Pt_0.4_ (P=0.004) remained associated with QoL independent of one another. After additional adjustment for positive, negative, and depressive symptoms, SCZ PRS (Pt_0.5_; P= 0.002) and wellbeing PRS (Pt_0.4_; P=0.04) remained associated with QoL. Moreover, only SCZ and subjective wellbeing PRSs were consistent with true polygenicity explaining a proportion of the variance in QoL, as may be appreciated by increasing degrees of explained variances and increasing significance levels with relaxing Pts (**Figure 2**).

As stated above, all final clinical phenotypes included in our regression model together explained 58.55% of the variability in QoL. By adding SCZ Pt_0.5_, MDD Pt_0.005_, and subjective wellbeing Pt_0.4_, the model explained 59.00% of the variability.

## Discussion

We here identified non-genetic factors contributing to QoL among patients suffering from SCZ. Our results show that up to 58% of variance in QoL may be explained using a range of demographic and clinical variables. We additionally demonstrate that genetic predisposition to SCZ and subjective wellbeing explain a (small) proportion of variability in QoL on top of clinical variables. The novelty of our method lies in the use of hypothesis-generating approaches to investigate a vast number of SCZ-associated genetic and non-genetic variables.

Most of the previous studies into QoL in SCZ had small to moderate sample sizes (Wang et al., 2017, Cruz et al., 2016, Savill et al., 2016) and have shown the association of particularly negative and positive symptoms with QoL in SCZ. A recent study in 157 SCZ patients showed the effects of excitement (Domenech et al., 2018), positive, negative, and depressive (Domenech et al., 2018) symptoms on QoL. The study by Domenech et al. used only clinical symptoms based on PANSS. We tested clinical symptoms assessed using multiple internationally well-established scales (e.g. CAPE, PANSS, CAN) and assessed a range of other phenotypic variables. Such rich phenotyping together with our large sample size allowed us to firmly establish additional variables associated with QoL in SCZ at increased statistical significance. Moreover, this approach allowed us to weigh the effect of all variables in one model. In line with our findings, a recent meta-analysis also found a substantial association between depressive symptoms and personal recovery, a concept related to quality of life (Van Eck et al., 2018).

Several of the variables we found to be associated with QoL in SCZ had to the best of our knowledge not been reported, such as disorganization, obsessive compulsive symptoms, suicidal attempts, unmet needs, acathisia, and cannabis craving. Although the underlying mechanisms of this latter association are still unclear, one may speculate that cannabis craving constitutes a proxy for cannabis abstinence, which in turn may increase anxiety and thus reduce psychological wellbeing. Alternatively, relatively high levels of cannabis dependence may worsen symptoms and thus negatively impact QoL.

Genetic predisposition to SCZ captured by PRS showed clear and persistent effects on QoL across all Pts. We also observed moderate effects of polygenic susceptibility to subjective well-being on QoL and no independent effects of genetic predisposition to MDD on QoL in our cohort.

The current study benefits from a large sample size of a multicenter prospective cohort study in the Netherlands with comprehensive phenotypic assessments in individuals with SCZ. The large sample size increases precision and reliability of our findings. The combination of a large sample size and rich phenotyping created a unique opportunity for a phenome-wide study to identify contributing factors to QoL in SCZ. In addition, carefully selected participants from several geographical locations restricted the risk of selection bias. On the other hand, several limitations should be borne in mind when interpreting our results. First, interpretation of principal component-driven variables may not be intuitive. Here, we managed to show its feasibility and usefulness. We reduced the number of variables of the four different domains of QoL into one variable and were able to assess the impact of multiple clinical and genetic determinants on this variable. Our results showed consistency in terms of direction and magnitude of the effect estimate when compared with ordinal domains of QoL (**Supplementary Figure 3**). Second, we are aware of relatively low power for genetic studies on a complex trait such as QoL both in our GWAS and LD score regression analyses. Our GWAS must therefore be regarded as a first exploratory GWAS of QoL in SCZ subjects, their siblings and healthy controls. Similarly, for LD score regression (LDSC), we were underpowered to reveal clear genetic correlations. LDSC analysis was done to explore possible genetic correlations with traits different from the ones we investigated and to investigate whether the trait with most significant genetic correlation results was identical to the trait with most significant PRS results, which indeed turned out to be the case. Third, in the current study population about 97% of participants were Caucasians which hampers generalizability to other ethnicities. Finally, our association analyses preclude us from drawing definite conclusions about causality. Future, well powered, prospective studies are necessary to improve insight into possible causal mechanisms.

In conclusion, we highlight multiple clinical and genetic associations with QoL that could be leveraged in daily care of patients with SCZ to improve their QoL. The variables highlighted in the current study could aid health professionals who interact with psychotic patients to more readily recognize the need for additional interventions in patients showing a high burden of such phenotypes. For example, although high levels of positive and negative symptoms are intuitively associated with QoL, disorganization, cannabis craving and obsessive-compulsive symptoms are also important contributors according to our analyses. Genetic risk scoring may furthermore be used to optimize identification of those SCZ patients susceptible to low quality of life, which in turn may advance timely management for these vulnerable patients.

## Supporting information

supplemental results

## Funding

The infrastructure for the GROUP study is funded through the Geestkracht programme of the Dutch Health Research Council (Zon-Mw, grant number 10-000-1001), and matching funds from participating pharmaceutical companies (Lundbeck, AstraZeneca, Eli Lilly, Janssen Cilag) and universities and mental health care organizations (Amsterdam: Academic Psychiatric Centre of the Academic Medical Center and the mental health institutions: GGZ Ingeest, Arkin, Dijk en Duin, GGZ Rivierduinen, Erasmus Medical Centre, GGZ Noord Holland Noord. Groningen: University Medical Center Groningen and the mental health institutions: Lentis, GGZ Friesland, GGZ Drenthe, Dimence, Mediant, GGNet Warnsveld, Yulius Dordrecht and Parnassia psycho-medical center The Hague. Maastricht: Maastricht University Medical Centre and the mental health institutions: GGZ Eindhoven en De Kempen, GGZ Breburg, GGZ Oost-Brabant, Vincent van Gogh voor Geestelijke Gezondheid, Mondriaan, Virenze riagg, Zuyderland GGZ, MET ggz, Universitair Centrum Sint-Jozef Kortenberg, CAPRI University of Antwerp, PC Ziekeren Sint-Truiden, PZ Sancta Maria Sint-Truiden, GGZ Overpelt, OPZ Rekem. Utrecht: University Medical Center Utrecht and the mental health institutions Altrecht, GGZ Centraal and Delta).

## Conflict of interest statement

All authors declare no conflict of interest.

