## supplemental results for "Phenome-wide and Genome-wide Analyses of Quality of Life in Schizophrenia"

### Supplemental Results - Polygenic risk scores are associated with quality of life in schizophrenia

**Supplementary Figure 1- Flowchart of the population for analysis.**

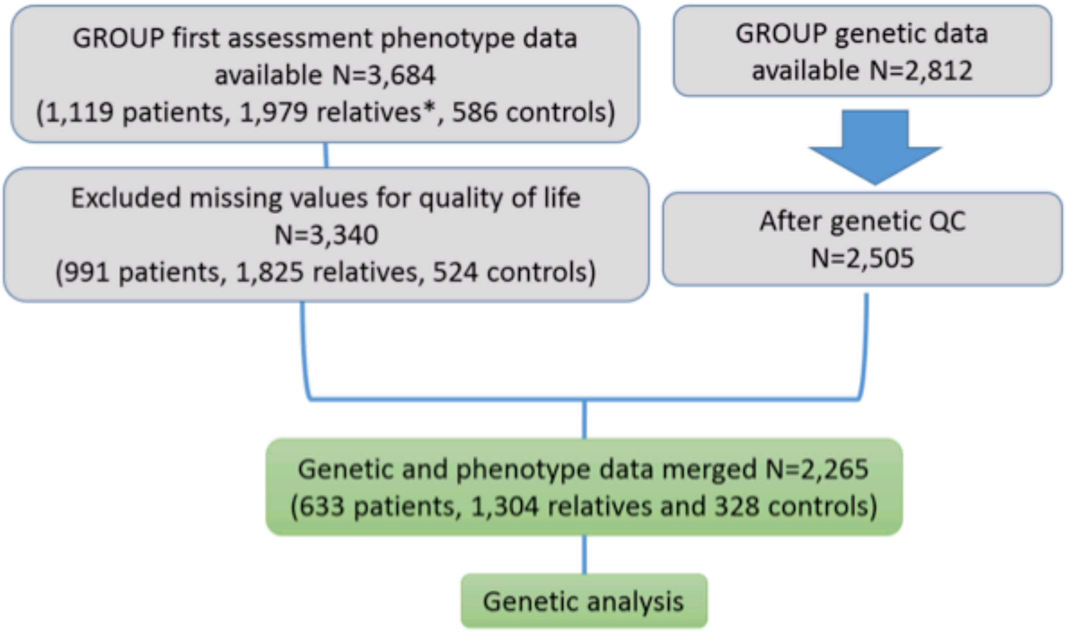


**Supplementary Figure 2- Heat map plot for the correlation of QoL domains and the principal components. The first principal component shows high degrees of correlation with all QoL domains.**

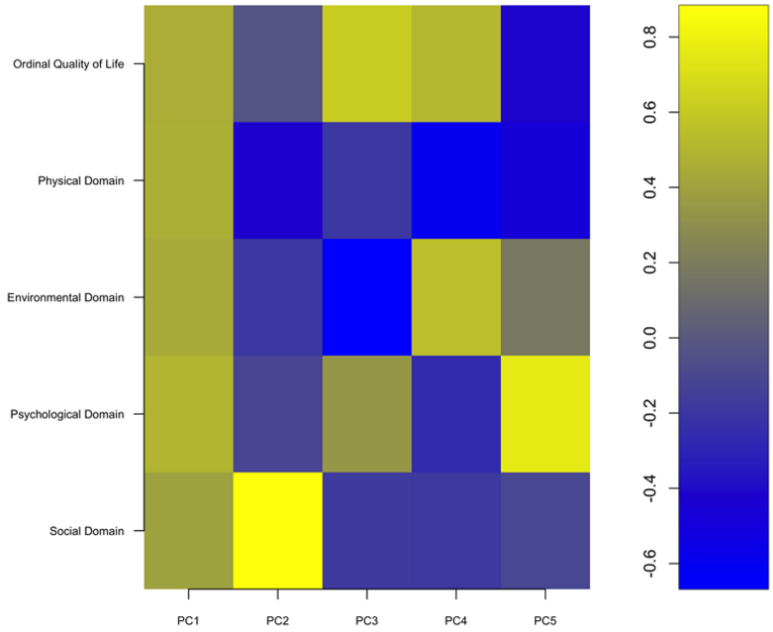

**Supplementary Figure 3- Volcano plot of the association test between clinical variables with QoL in SCZ.**

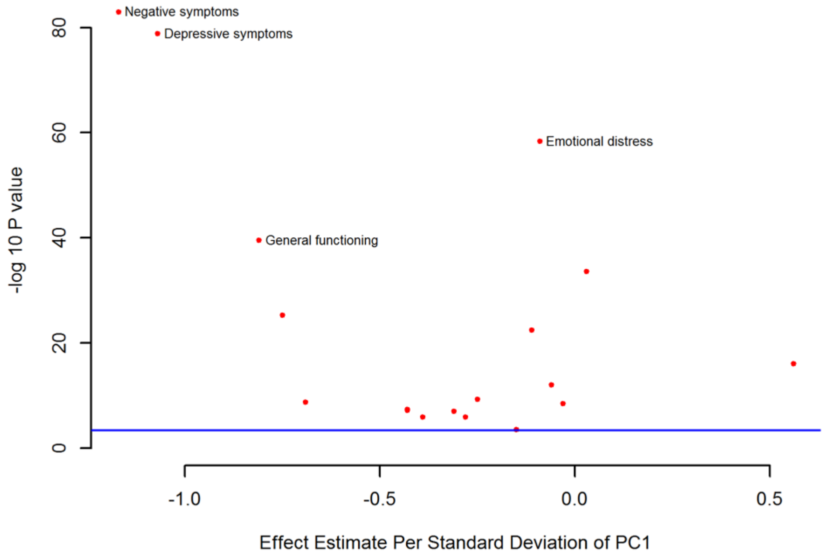

**Supplementary Figure 4-Box plot for the association of negative and depressive symptoms with ordinal domain of QoL.**

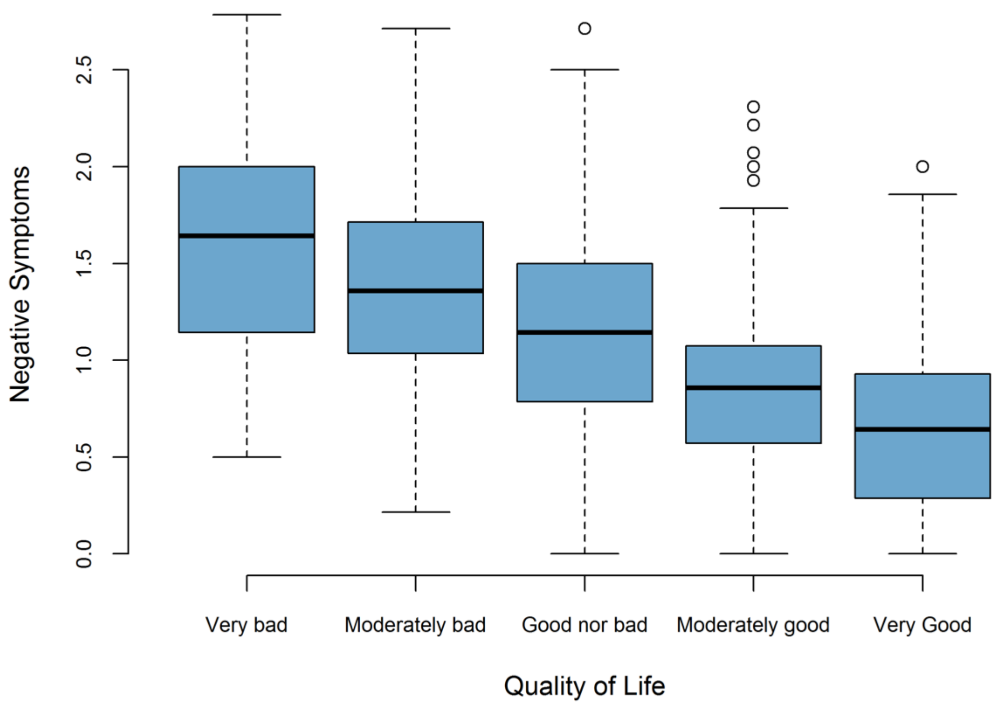


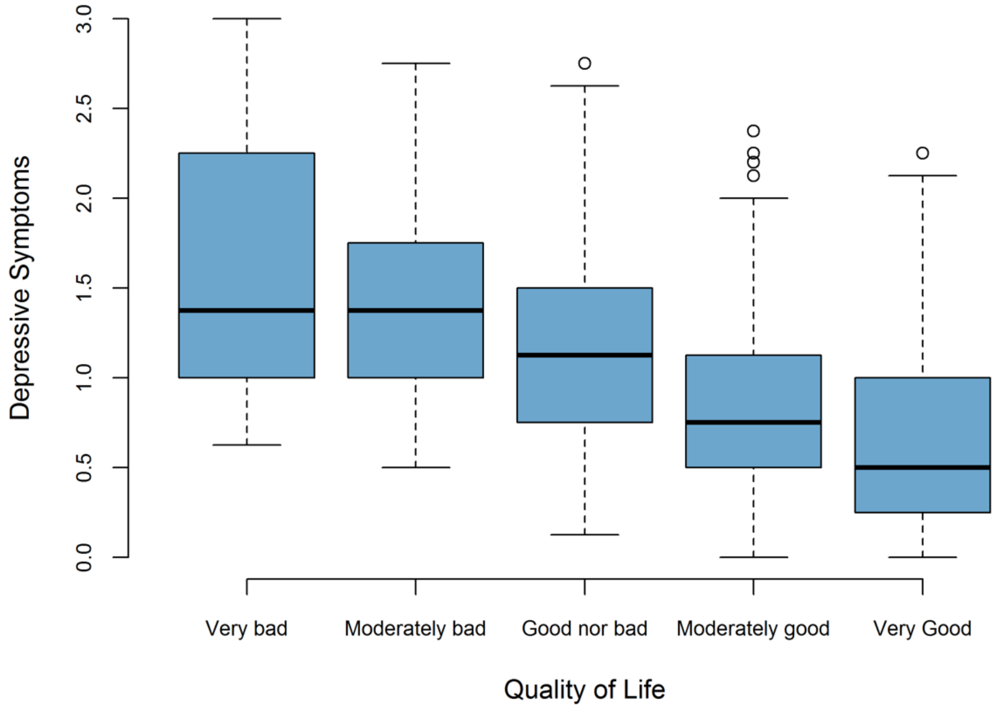

**Supplementary Figure 5- PRS analysis for MDD vs. QoL.** The figure illustrates the results using linear mixed models. Displayed are the strengths of the association results (-log 10 P) and explained variances per Pt (p-value threshold).

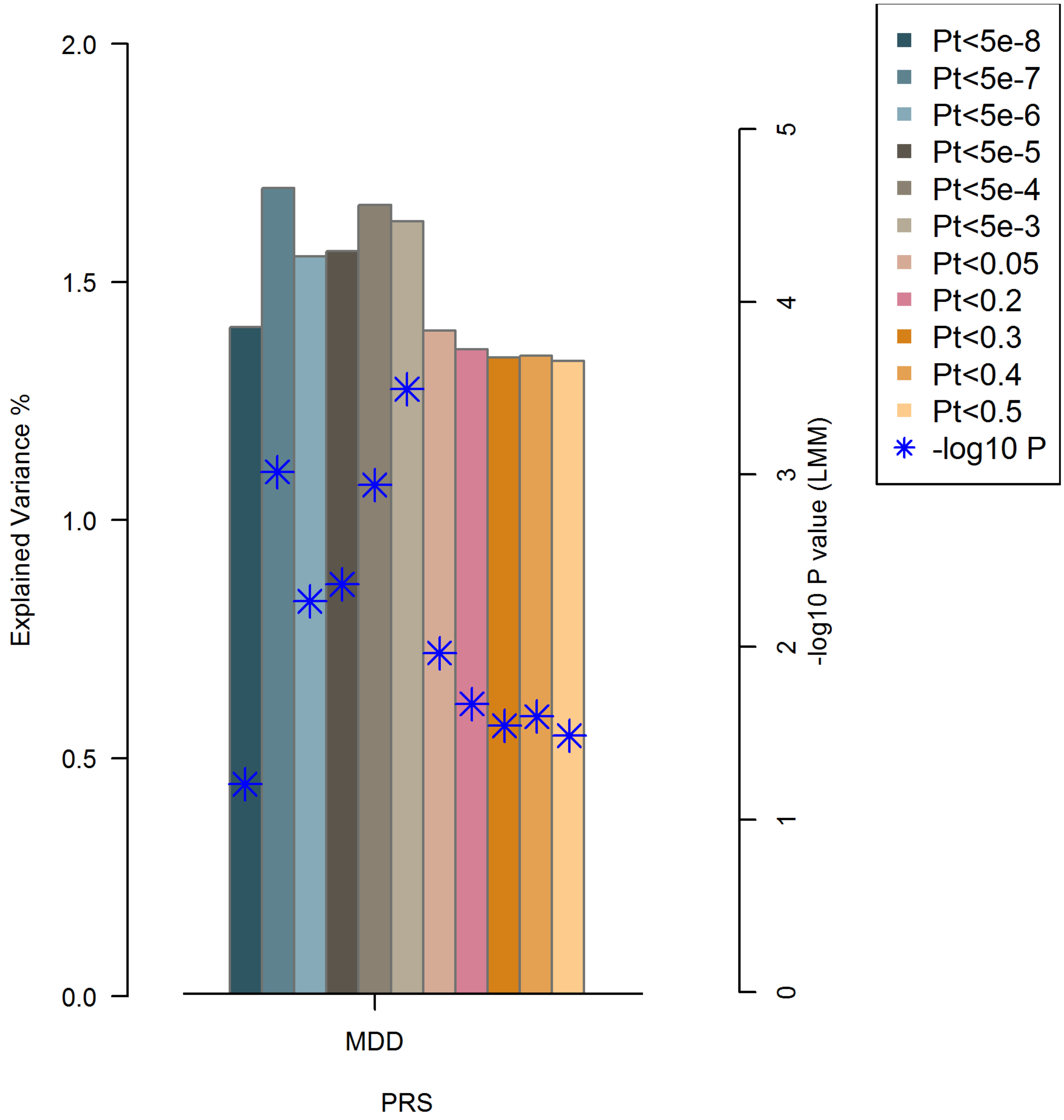

**Supplementary Figure 6 – Schizophrenia PRS analysis in patients only (N=633).** This figure demonstrates findings for the patients only subgroup are similar to the entire study population (patients, relatives and healthy controls), albeit at lower significance which is expected given the smaller sample size, but with the same P-value threshold of strongest associations at Pt=0.5 ('0p5' in the graph).

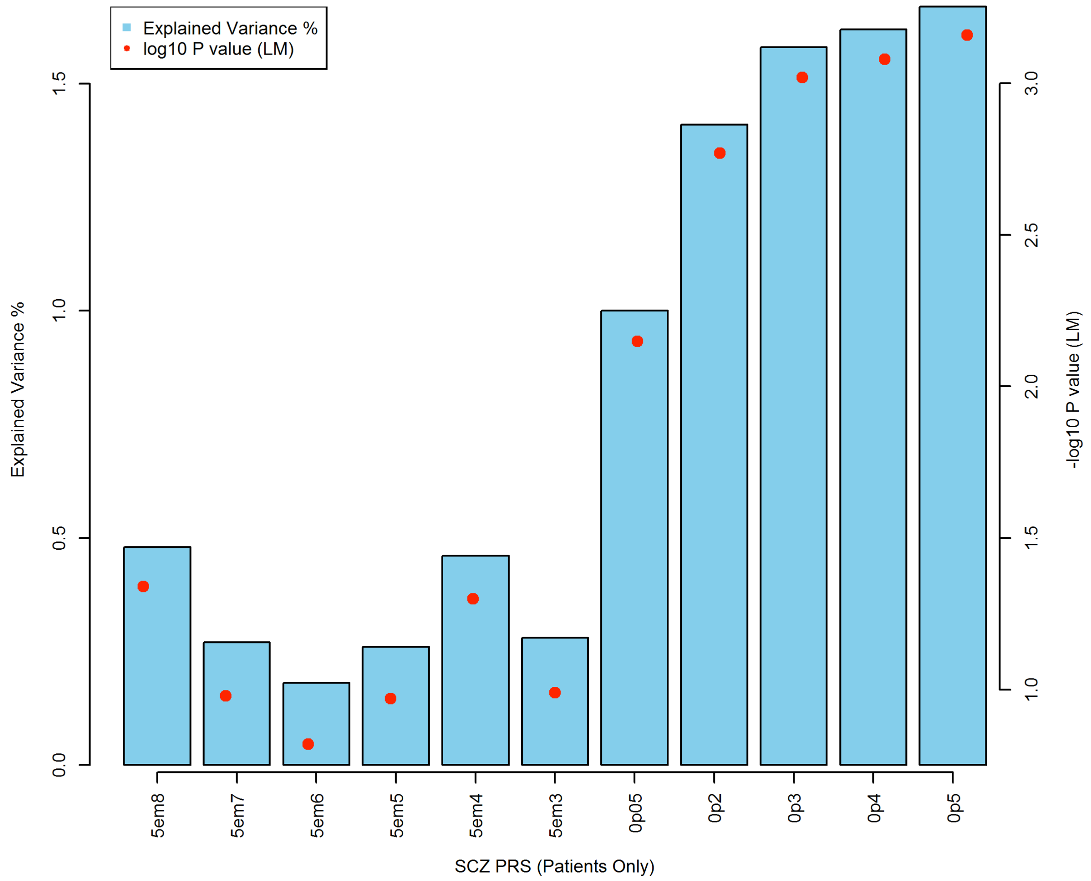

**Supplementary Figure 7**

The primary BOLT-LMM GWAS of QoL in the whole dataset after QC (N=2,265) and all three sensitivity analyses identified no locus surpassing the genome-wide significance threshold ($5\times10^{-8}$; **Supplementary Figure 7)**. At $p < 10^{-5}$ in the primary BOLT-LMM GWAS, seven independent loci were associated with QoL, in the vicinity of *CYP4F3, GNB1L, UBXN2B, GRIA1, THBD, CLSTN2-AS1, LOC100506272* genes (**Supplementary Figure 8)**.

| SNP | CHR | BP | EA | AA | EAF | BETA | SE | P value | Gene |
|---|---|---|---|---|---|---|---|---|---|
| **rs4808352** | 19 | 15777931 | C | T | 0.226269 | 0.178919 | 0.0362702 | 8.1×10-07 | *CYP4F3* |
| **rs2301558** | 22 | 19751829 | T | C | 0.254746 | -0.164087 | 0.0348483 | 2.5×10-06 | *GNB1L* |
| **rs12681674** | 8 | 59297629 | G | A | 0.120088 | 0.213345 | 0.0463611 | 4.2×10-06 | *UBXN2B* |
| **rs72804672** | 5 | 153242887 | C | T | 0.139514 | 0.201893 | 0.0439584 | 4.4×10-06 | *GRIA1* |
| **rs2424506** | 20 | 23036651 | A | G | 0.428918 | 0.137637 | 0.0301564 | 5.0×10-06 | *THBD* |
| **rs74876817** | 3 | 140264008 | T | C | 0.0311258 | -0.389057 | 0.085462 | 5.3×10-06 | *CLSTN2-AS1* |
| **rs115948119** | 4 | 188468685 | C | T | 0.0103753 | -0.672053 | 0.149383 | 6.8×10-06 | *LOC100506272* |

CHR: Chromosome; BP: position; EA: Effect Allele; AA: Alternative Allele; EAF: effect allele frequency; SE: Standard Error; Gene: the gene located nearest to the associated SNP.

a) LMM BOLT primary GWAS of QoL in 2,265 patients, relatives and controls: Manhattan plot and Q-Q plot (lambda =1.01).

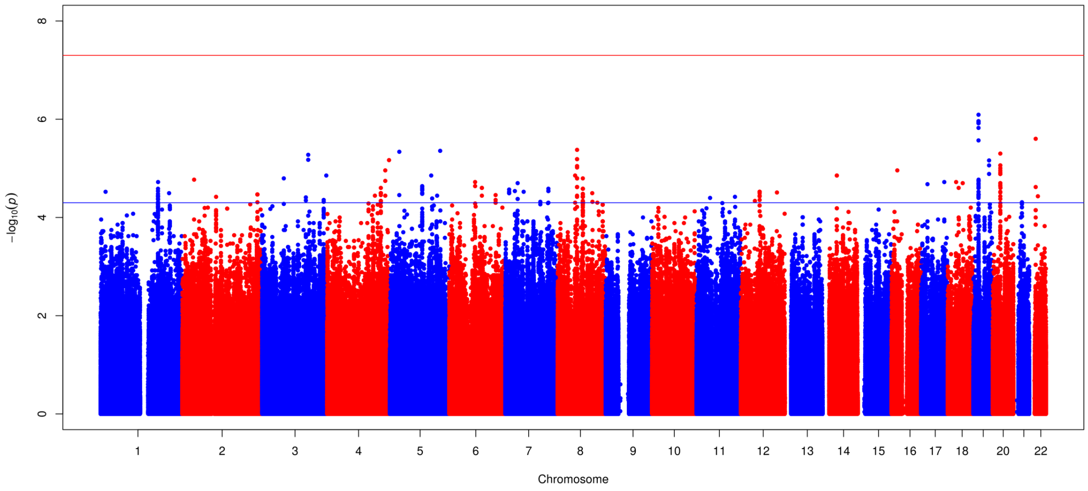


a) QQ Plot of LMM BOLT GWAS

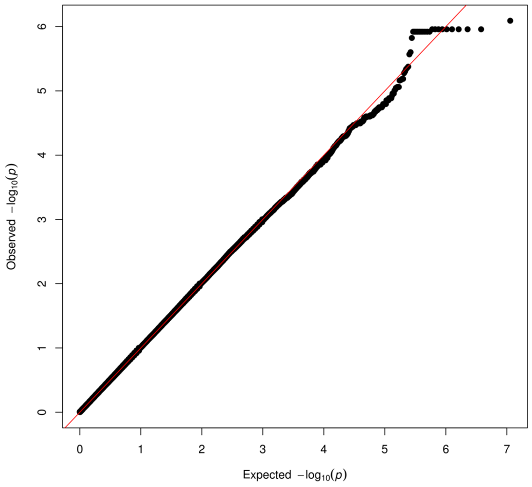

B. Manhattan plot and QQ plot (lambda = 1.019) of the first sensitivity analysis: GWAS of QoL in 1,069 unrelated subjects (after randomly selecting one individual from each family; including 495 schizophrenia patients).

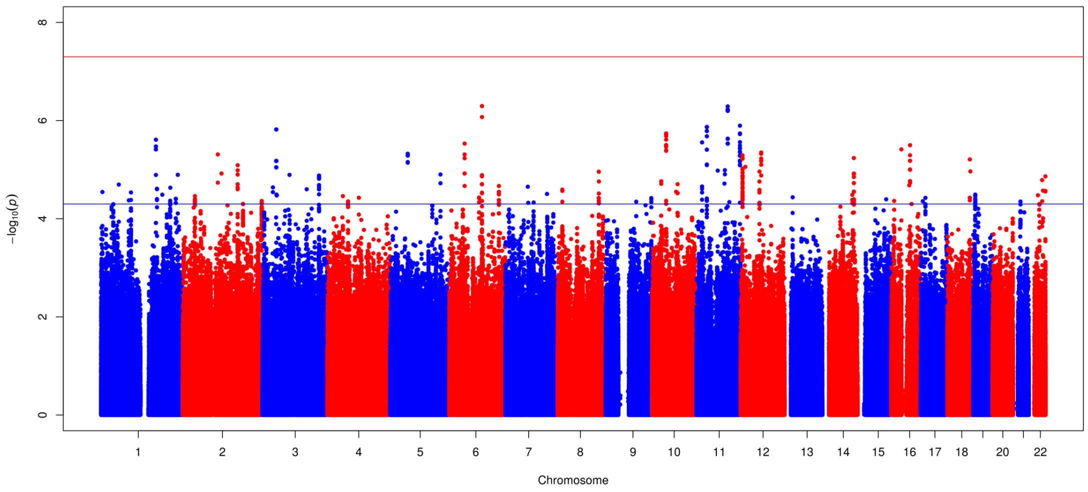


Note: Blue line is suggestive significance threshold (p=5X10$^{-5}$), and red line is genome-wide significance threshold (p=5X10$^{-8}$).

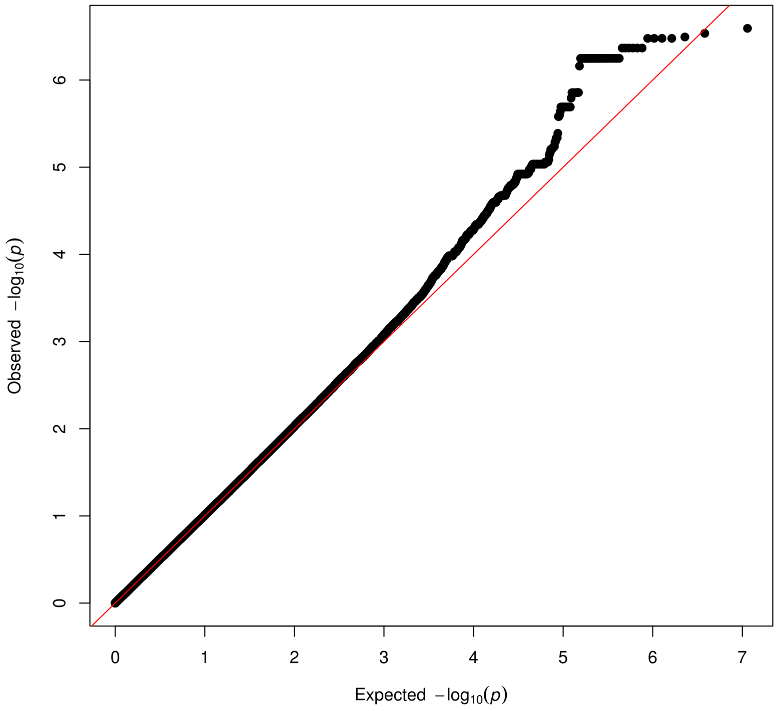

C. Manhattan plot and QQ plot (lambda = 1.015) of the second sensitivity analysis in schizophrenia patients only (N= 605 unrelated SCZ patients).

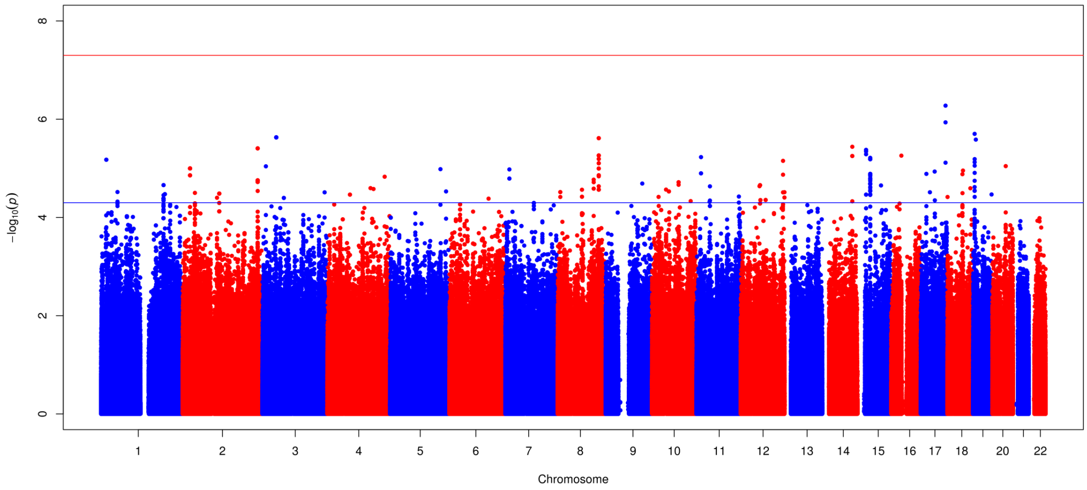


Note: Blue line is suggestive significant threshold ($p=5X10^{-5}$), and red line is genome-wide significant threshold ($p=5X10^{-8}$).

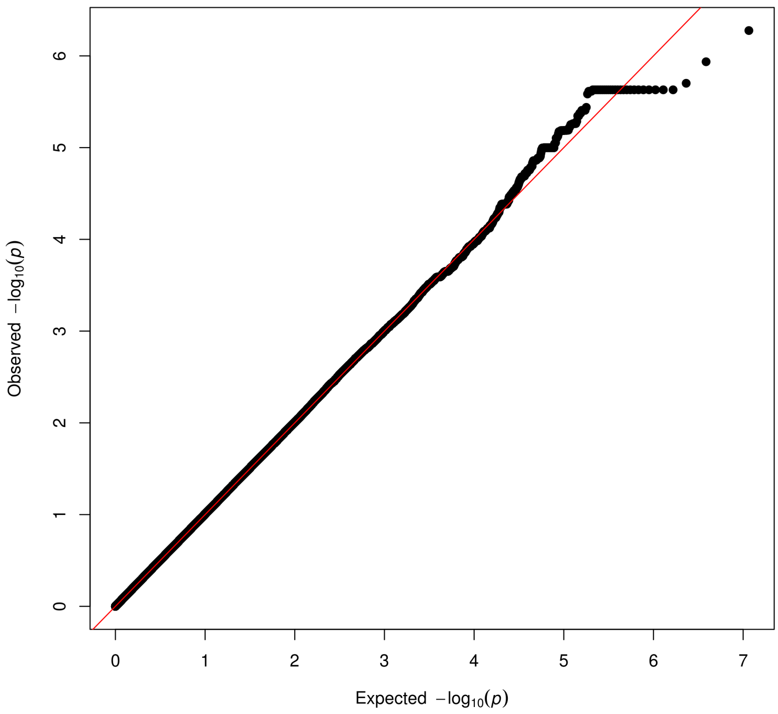

D. Manhattan plot and QQ plot (lambda = 1.013) of the third sensitivity analysis (only healthy subjects): GWAS of QoL in 953 unrelated healthy people (by preferably selecting parent pairs (i.e. two parents from the same family) and if those are unavailable randomly selecting one healthy subject within a family).

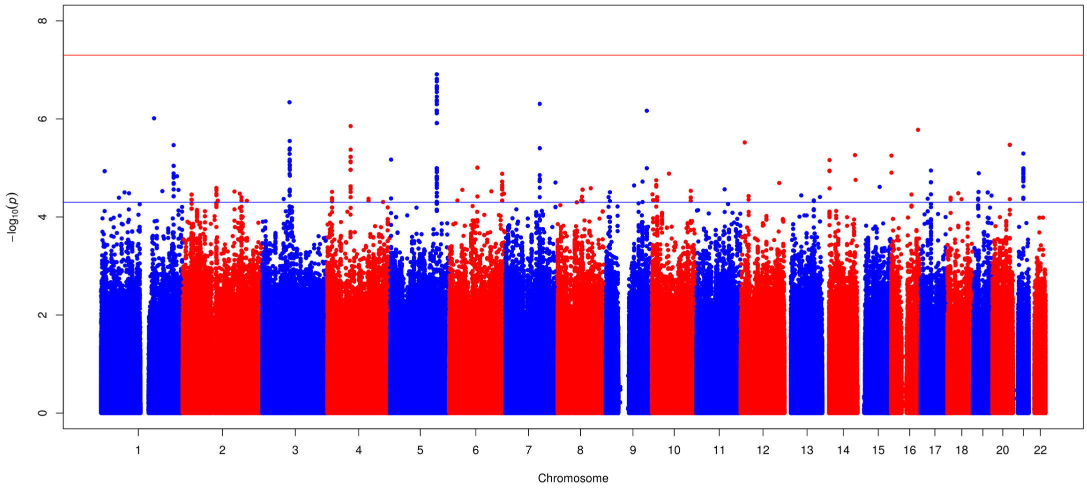


Note: Blue line is suggestive significant threshold ($p=5X10^{-5}$), and red line is genome-wide significance threshold ($p=5X10^{-8}$).

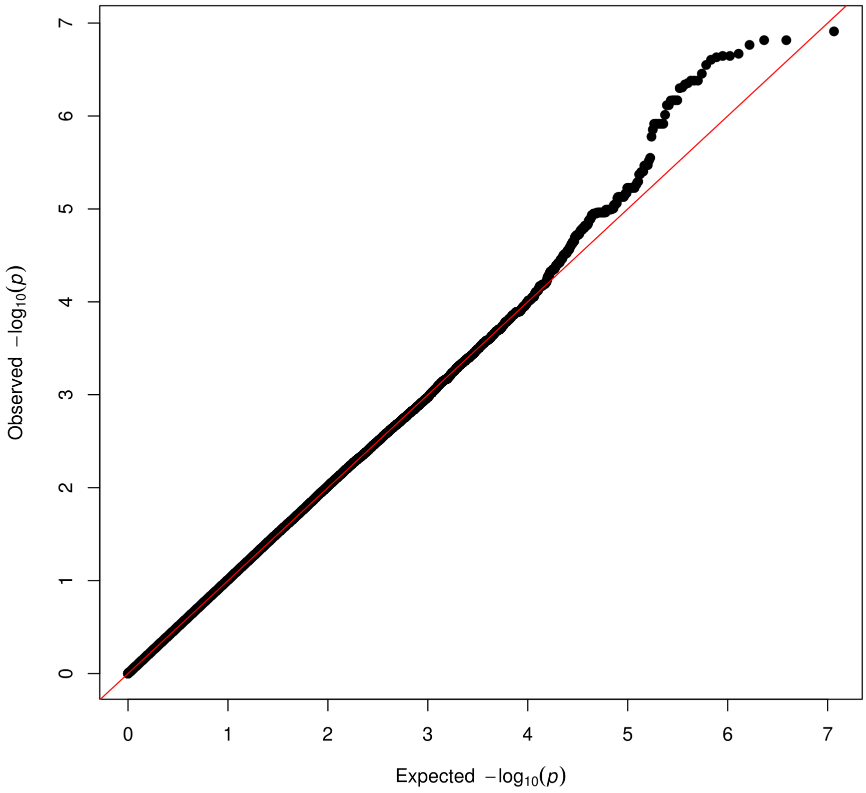

**Supplementary Figure 8. Locus Zoom plots for LMM BOLT primary GWAS most significant signals.**

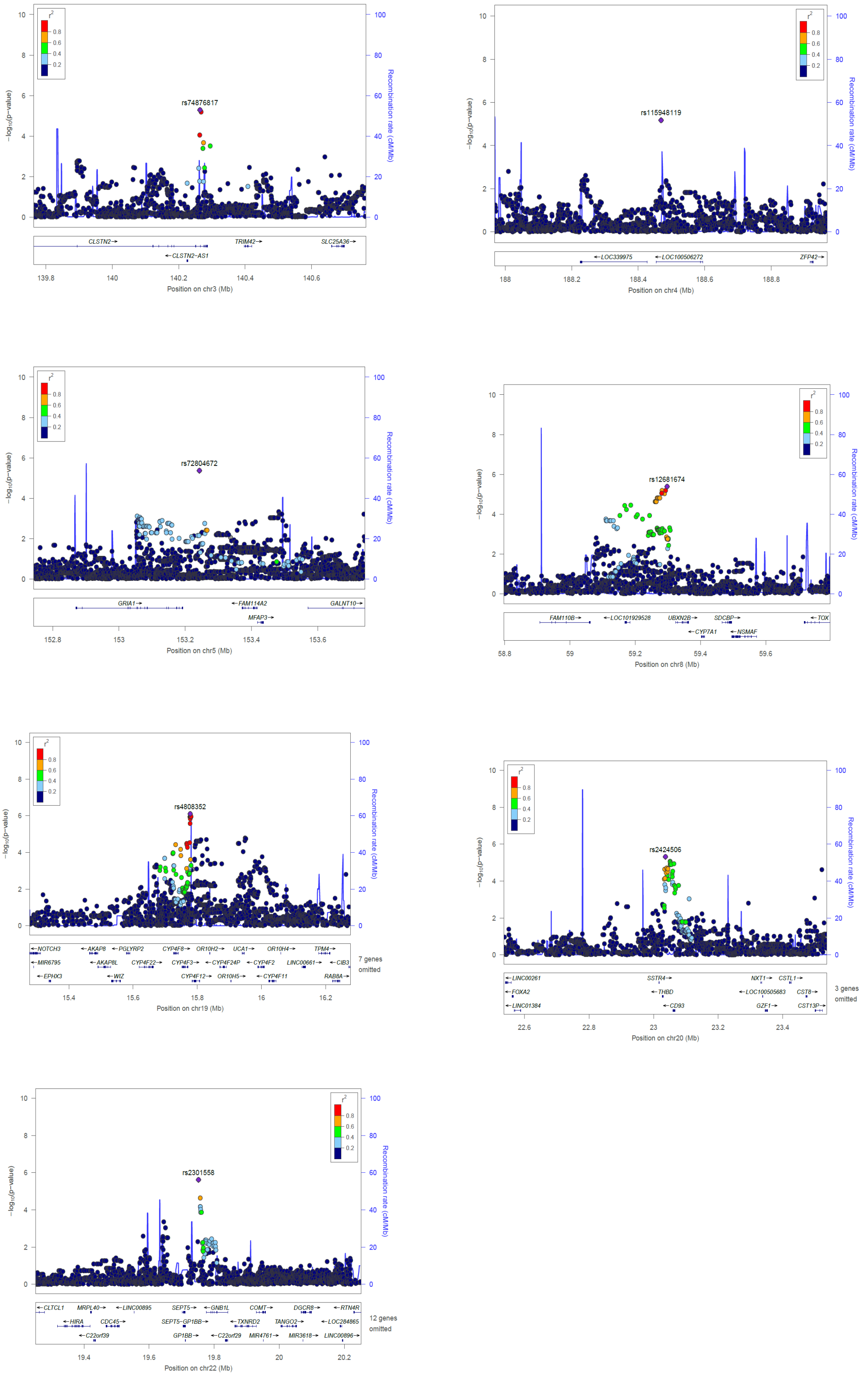

**Supplementary Table 1. The complex-LD regions (build GRCh37) were removed before PRS calculation.**

| Chromosome | Base pair start | Base pair end |
|---|---|---|
| 6 | 25392021 | 33392022 |
| 8 | 111930824 | 114930824 |
| 11 | 46043424 | 57243424 |
| 1 | 48287980 | 52287979 |
| 2 | 86088342 | 101041482 |
| 2 | 134666268 | 138166268 |
| 2 | 183174494 | 190174494 |
| 3 | 47524996 | 50024996 |
| 3 | 83417310 | 86917310 |
| 3 | 88917310 | 96017310 |
| 5 | 44464243 | 50464243 |
| 5 | 97972100 | 100472101 |
| 5 | 128972101 | 131972101 |
| 5 | 135472101 | 138472101 |
| 6 | 56892041 | 63942041 |
| 6 | 139958307 | 142458307 |
| 7 | 55225791 | 66555850 |
| 8 | 7962590 | 11962591 |
| 8 | 42880843 | 49837447 |
| 10 | 36959994 | 43679994 |
| 11 | 87860352 | 90860352 |
| 12 | 33108733 | 41713733 |
| 12 | 111037280 | 113537280 |
| 20 | 32536339 | 35066586 |

**Supplementary Table 2. Strongest genetic correlation results for quality of life using LD Hub, ordered by P value.**

| trait | PMID | Category | ethnicity | rg | se | z | p | h2_obs | h2_obs_se | h2_int | h2_int_se | gcov_int | gcov_int_se |
|---|---|---|---|---|---|---|---|---|---|---|---|---|---|
| Schizophrenia | 25056061 | psychiatric | Mixed | 0.4917 | 0.2974 | 1.6536 | 0.0982 | 0.4593 | 0.0194 | 1.0537 | 0.0141 | 0.0359 | 0.0079 |
| Subjective well being | 27089181 | psychiatric | European | -0.5961 | 0.4020 | -1.4830 | 0.1381 | 0.0247 | 0.0022 | 1.0042 | 0.0081 | 0.0005 | 0.0053 |
| Years of schooling (proxy cognitive performance) | 25201988 | education | European | -0.3181 | 0.2207 | -1.4415 | 0.1494 | 0.1088 | 0.0080 | 1.0225 | 0.0102 | 0.0032 | 0.0057 |
| FEV1/FVC* | 28166213 | lung_function | European | -0.5054 | 0.3560 | -1.4198 | 0.1557 | 0.2569 | 0.0215 | 0.9747 | 0.0105 | 0.0040 | 0.0056 |
| Multiple sclerosis | 21833088 | autoimmune | European | 0.5831 | 0.4239 | 1.3755 | 0.1690 | 0.0555 | 0.0302 | 1.0596 | 0.0104 | -0.0120 | 0.0063 |
| PGC cross-disorder analysis | 23453885 | psychiatric | European | 0.4001 | 0.3060 | 1.3077 | 0.1910 | 0.1741 | 0.0127 | 1.0118 | 0.0113 | 0.0306 | 0.0070 |
| Major depressive disorder | 22472876 | psychiatric | European | 0.4771 | 0.5731 | 0.8325 | 0.4051 | 0.164 | 0.0288 | 1.0081 | 0.0071 | 0.0074 | 0.005 |

Note: rg: the estimated genetic correlation; se: the bootstrap standard error of the genetic correlation estimate; z: the z-statistics; p: p-value from the z-statistic; h2_obs: estimated snp-heritability of the second phenotype; h2_obs_se: bootstrap standard error of the snp-hertiability estimate; h2_int: LD score regression intercept for the second phenotype; h2_int_se: bootstrap standard error of the of the intercept; gcov_int: estimated genetic covariance between p1 and p2, if there is no sample overlap, the gcov_int will near zero; gcov_int_se: bootstrap standard error of the genetic covariance. FEV1/FVC: Forced expiratory volume in 1 second (FEV1)/Forced Vital capacity (FVC).

**Supplementary Table 3- The clinical variables independently associated with QoL in the backward stepwise linear model (N= 925 schizophrenia patients).**

| Variable | Scale | Standardized Effect Estimate | Standard Error | P Value |
|---|---|---|---|---|
| Intercept | NA | 1.14 | 0.22 | $2\times10^{-7}$ |
| Emotional distress, points | PANSS | -0.04 | 0.007 | $1\times10^{-9}$ |
| Cannabis craving, yes | OC-DUS | -0.15 | 0.06 | 0.01 |
| Remission status, yes | PANSS | 0.15 | 0.07 | 0.03 |
| sex (female) | NA | 0.10 | 0.07 | 0.13 |
| Number of unmet needs, points | CAN | -0.23 | 0.09 | 0.02 |
| Positive symptoms, points | CAPE | 0.26 | 0.07 | $2\times10^{-4}$ |
| Excitement, points | PANSS | 0.02 | 0.01 | 0.03 |
| Depressive symptoms, points | CAPE | -0.43 | 0.08 | $6\times10^{-8}$ |
| Global assessment of functioning (disabilities), points* | GAF | 0.01 | 0.002 | $2\times10^{-10}$ |
| age | NA | -0.01 | 0.003 | $3\times10^{-3}$ |
| Negative symptoms, points | CAPE | -0.68 | 0.07 | $2\times10^{-16}$ |

CAPE: Community Assessment of Psychic Experiences; PANSS: positive and negative syndrome scale; GAF, Global assessment of functioning; CAN, The Camberwell Assessment of Need; OC-DUS: obsessive-compulsive drug use scale; * Greater score indicates better functioning.
