## supplemental methods for "Phenome-wide and Genome-wide Analyses of Quality of Life in Schizophrenia"

### Supplemental Methods - Polygenic risk scores are associated with quality of life in schizophrenia

*Quality of Life: Computation of a single principal component*

The WHOQOL-BREF scale includes 24 items covering four domains of QoL (1), namely physical health (seven items), psychological (six items), social relations (three items), and environment (eight items). The questionnaire additionally provides a general estimate of QoL presented as 5 categories "very bad", "moderately bad", "good nor bad", "moderately good", and "very good". To reduce the number of QoL variables and create a suitable variable for statistical analyses, we performed principal component analysis to preferably identify one quantitative component that best explained most of the variability in QoL domains. Similar to a previously published GWAS using principal component analysis to derive a single phenotype (2), the percentage of variance accounted for by the first unrotated principal component was computed. This principal component was standardized for further analysis, explained 62.5% of the variability in our data and correlated well with all WHOQOL-BREF domains (**Supplementary Figure 2**) whereas the other four components were correlated with only one of the domains and showed minimal correlations with other domains of QoL. We therefore used this first principal component for subsequent analyses.

*Genotype data and GWAS of QoL*

Genotype data for 2,812 GROUP participants was generated on a customized Illumina IPMCN array with 570,038 single nucleotide polymorphisms (SNPs). This chip contains ~250k common SNPs, 250K Exome chip variants (rare, exonic, nonsynonymous, MAF < 1%), and ~50K psychiatric-related variants. Quality control procedures were performed using PLINK v1.9(3). SNPs and samples with call rates below 95% and 98%, respectively, were removed. A strict SNP QC only for subsequent sample quality control steps was conducted. This involved a minor allele frequency (MAF) threshold>10% and a Hardy-Weinberg equilibrium

(HWE) P-value>$1\times10^{-05}$, followed by linkage disequilibrium (LD) based SNP pruning ($R^2$<0.2). This resulted in ~58K SNPs to assess sex errors, heterozygosity (F<3 standard deviation (SD)), homozygosity (F>3SD) and relatedness by pairwise identity by descent (IBD) values. Duplicate samples (pihat > 0.8) were removed and remaining pairs were manually checked since this dataset contains family members. After removing failing samples (n=208), a regular SNP QC was performed (SNP call rate>98%, HWE P-value >$1\times10^{-06}$, MAF>1%). After multi-dimensional scaling (MDS) clustering with Hapmap Phase 3 individuals to check ethnicity, samples that deviated more than 3 standard deviations from our dataset were removed (n=91). Next, strand ambiguous SNPs and duplicate SNPs were removed. Mendelian errors were set to missing followed by another missingness check (2% threshold) for samples (n=8) and SNPs, and SNPs with a differential missingness between cases and controls were removed. In total, 2,505 individuals and 275,021 SNPs passed these abovementioned QC steps. After merging with the phenotype file, 2,265 individuals were left for genetic analyses (**Supplementary Figure 1**).

Additional SNPs were imputed on the Michigan server(4) using the HRC r1.1 2016 reference panel with European samples after phasing with Eagle v2.3. Post-imputation QC involved removing SNPs with an imputation quality score<0.3, with a MAF<0.01, SNPs that had a discordant MAF compared to the reference panel, and strand ambiguous AT/CG SNPs and multi-allelic SNPs.

We performed linear mixed models (LMM) association testing implemented in BOLT-LMM (v2.3) software(5) to assess associations between SNPs and QoL (**supplementary information**). BOLT-LMM corrects for confounding from population structure and cryptic relatedness. We assumed additive genetic models on the continuous first principal component of QoL adjusted for age, age$^2$, sex, and the first three genetic principal components. We used the generally accepted association P-value threshold of $P < 5\times10^{-8}$ for genome-wide significance. To account for population stratification, we corrected P values for genomic inflation value (lambda), (Supplementary results, Supplementary Figure 7 A & 8) . As sensitivity analyses, we performed GWASs of unrelated individuals within 3 separate unrelated groups. Cohort 1 (Supplementary results, Supplementary Figure 7B) included unrelated individuals in the whole sample (selected by random selection of one individual from each family; N=1,069, consisting of 495 SCZ patients, 180 healthy sibs, 130 healthy parents and 291 controls). Cohort 2 (Supplementary results, Supplementary Figure 7C) included only unrelated SCZ patients by random selection of one SCZ patient from each family (N=605). Cohort 3 (Supplementary resultd, Supplementary Figure 7D) included unrelated participants not suffering from SCZ (so only healthy unrelated subjects) by preferably selecting parent pairs (i.e. two parents from the same family) and if those were unavailable randomly selecting one healthy subject within a family (N=953, consisting of 410 healthy parents, 252 healthy sibs and 291 healthy controls). To verify the subjects in all these cohorts were indeed unrelated, IBDs in all 3 cohorts were calculated using genome-wide autosomal SNPs. As all IBD pi-hats turned out to be < 0.1, all these selected subjects in the three cohorts were included in the three sensitivity analyses. These sensitivity GWASs were conducted using linear models implemented in Plink with the same covariates as the primary BOLT-LMM analysis.

*PRS analyses*

We calculated explained variance and P values for PRSs on QoL in two stages. In the first stage, we analyzed the effect of age, sex and disease status on QoL to obtain residuals of this model. Subsequently, we tested the association of the residuals of the previous model with each of the various PRSs using a linear mixed model (LMM) with family ID as random effect and the first three genetic principal components as covariates. We additionally included the most significantly associated PRSs from each SCZ, MDD, and wellbeing PRS group in one single mixed model to assess if these PRSs were statistically independent of each other. We subsequently included negative, positive, and depressive symptoms as covariates (as these were previously known to associate with QoL, which was also confirmed in the current cohort), and PRSs in mixed models to assess the additional explained variance of the three PRSs beyond clinical phenotypes. We additionally performed sensitivity SCZ PRS analyses on patients only to assess whether we observed similar effects to the combined set of patients, siblings and controls.

**Appendix 1**- Demographic, symptom-level, family loading, social cognition, IQ, medication use, and theory of mind phenotypes included in this study.

| Phenotypes | |
|---|---|
| Demographic | Gender |
| | Non-white ethnic minority status (y/n) |
| | Member of multiple birth (y/n) |
| | Medical Questionnaire: Does subject have a twin? (y/n) |
| | Region where a patient lives |
| | Weight at birth (Composite file) |
| | Work, fulltime or part-time (Composite file) |
| | Work, paid or voluntary (Composite file) |
| | Work-related activities Composite file) |
| | Year of birth |
| Clinical | Age of Onset first Psychosis |
| | Amphetamine positive by urinalysis (yes or no) |
| | Duration of Illness (years) |
| | Duration untreated psychosis to first contact with mental health care (years) |
| | GAF (Global Assessment of Functioning), disabilities |
| | GAF (Global Assessment of Functioning), symptoms |
| | BARS (Barnes Akathisia Rating Scale Global), clinical assessment of Akathisia |
| | Recent Onset Psychosis - 12 months |
| | Recent Onset Psychosis - 24 months |
| | SDS (Schedule for the Deficit Syndrome), Fulfils criteria of deficit syndrome (y/n) |
| | Sort Antipsychotics oral (first or second generation) |
| | Sort Antipsychotics depot (first or second generation) |
| | Status Antipsychotics use (yes or no) |
| | Status of using other non-antipsychotic medication (yes or no) (Composite file) |
| | UPDRS (Unified Parkinson's Disease Rating Scale) overall score |
| Composite International Diagnostic Interview (CIDI) | CIDI: 12 month any cannabis use; Alcohol number of units per week; Cannabis 12 months most intensive mode of use |
| | (CIDI) Cocaine use ever (y/n) |
| | Cocaine positive by urinalysis (y/n) |
| | CIDI Coke use (frequency, lifetime) |
| | CIDI Coke use (lifetime, y/n) |
| | CIDI, other drug use (frequency, lifetime) |
| | CIDI, other drug use (y/n, lifetime) |
| | CIDI PCP (Phenylcyclidine) use (frequency, lifetime) |
| | CIDI PCP (Phenylcyclidine) use (y/n, lifetime)<br>Positive scale (PANSS, Positive and Negative Syndrome Scale)<br>Positive symptoms (PANSS, Positive and Negative Syndrome Scale) |
| | CIDI Psychedelics use (frequency, lifetime) |
| | CIDI Psychedelics use (yes or no, lifetime) |
| | CIDI Stimulants use (frequency, lifetime) |
| | CIDI Stimulants use (yes or no, lifetime) |
| Community Assessment of Psychic Experiences (CAPE) | CAPE depressive symptoms, distress |
| | CAPE Depressive symptoms, frequency |
| | CAPE Negative symptoms, distress |
| | CAPE Negative symptoms, frequency |
| | CAPE Positive symptoms, distress |
| | CAPE Positive symptoms, frequency |
| Word Learning Task (WLT) | 15WLT delayed recall items correct |
| | 15WLT recognition, number of correct negatives |
| | 15WLT recognition, number of correct positives (hits) |
| | 15WLT recognition, total correct |
| | 15WLT, retention rate |
| | 15WLT total items correct, immediate recall |
| Abnormal Involuntary Movement Scale (AIMS) | AIMS, overall score |
| | AIMS, Dystonia (y/n) |
| | Average moves between 10-14yrs (x 100) |
| | Average moves between 15-19yrs (x 100) |
| | Average moves between 20-39yrs (x 100) |
| | Average moves between 40-59yrs (x 100) |
| | Average moves between 5-9yrs (x 100) |
| Continuous Performance Test (CPT-HQ) | accuracy, overall |
| | RT correct responses |

| | |
|---|---|
| | number of correct positives (hits) |
| | number of false negatives |
| | number of false positives |
| | sensitivity index ((hits - false positive target)/28) * 100 |
| Degraded facial affect recognition task (DFAR) | percentage angry faces |
| | percentage fearful faces |
| | percentage happy faces |
| | percentage neutral faces |
| | percentage total correct |
| | total items correct |
| Premorbid Adjustment Scale (PAS) | PAS mean age 12-16 (Personality Assessment Screener) |
| | PAS mean age 16-19 (Personality Assessment Screener) |
| | PAS mean up to age 12 (Personality Assessment Screener) |
| | PAS overall score (Personality Assessment Screener) |
| | Premorbid Adjustment Scale (PAS), Disorganization |
| | Premorbid Adjustment Scale (PAS) Emotional distress |
| | Premorbid Adjustment Scale (PAS), Excitement |
| Camberwell Assessment of Need (CAN) | CAN measures the met and unmet needs of subjects |
| Comprehensive Assessment of Symptoms and History (CASH) | CASH, Education, subject completed |
| | CASH, Educational degree, highest |
| Schedules Clinical Assessment Neuropsychiatry (SCAN) | CASH or SCAN Diagnosis DSM-IV axis I |
| | CASH or SCAN Differential diagnosis |
| | CASH or SCAN Differential diagnosis DSM-IV axis I |
| Composite file (a file designed for the purpose of the GROUP study, consisting of several questions) | Average moves between 0-4yrs (x 100), urbanicity |
| | Marital Status (married y/n) |
| | Ethnicity Subject |
| | Ethnicity, country of origin grandfather/ grandmother |
| | Current household |
| | Current marital status |
| | Daily Dose Antipsychotics 1 |
| | Name Antipsychotics 1 |
| | Daily Dose Antipsychotics 2 (in the event another antipsychotic was taken) |
| | Environmental factors, number of moves |
| | Environmental factors, subject has attempted suicide |
| | Environmental factors, subject has children |
| | Environmental factors, subject has had (a) staying back(s) |
| | Environmental factors, subject has lost parent |
| | Environmental factors, subject has lost sibling |
| | number of moves (lifetime) |
| | number of Psychotic Episodes |
| Wechsler Adult Intelligence Scale (WAIS) | block design raw score |
| | block design scaled score |
| | calculation raw score |
| | calculation scaled score |
| | digit symbol substitution raw score |
| | digit symbol substitution scaled score |
| | Estimation total IQ |
| | information raw score |
| | information scaled score |
| | total scaled scores |
| Yale-Brown Obsessive Compulsive Scale (YBOCS) | Overall score |
| | last week obsessions or compulsions |
| | Resistance to Obsessions and Compulsions |
| | Severity Compulsions |
| | Severity Obsessions |
| | Total score |
| Family Interview for Genetic Studies (FIGS) | Family loading bipolar disorder |
| | Family loading drug abuse |
| | Family loading psychotic disorder |
| | number of Relatives |
| Response Shifting Task (RST) | accuracy cost score (rai-rar) |
| | proportional cost index score ((rai-rar)/rar) |
| | reaction time cost index score ((rtr-rti)/rti) |
| | reaction time cost score (rtr-rti) |
| Positive and Negative Syndrome Scale (PANSS) | General Psychopathology scale |
| | Negative scale |
| | Negative symptoms |
| | PANSS Remission Subject in remission? (cross-sectional) |
| Obsessive-compulsive drug use scale (OCD-DUS) | version cannabis OC Cannabis craving |
| | version cannabis C Cannabis thoughts |

| | |
|---|---|
| | version cannabis OC Total score |
| Hints (theory of mind task) | Hints sensitivity |
| | Hints total score |
